# Inferring B-cell derived T-cell receptor induced multi epitope-based vaccine candidate against enterovirus 71 (EV 71): A reverse vaccinology approach

**DOI:** 10.1101/2023.03.04.531076

**Authors:** Subrat Kumar Swain, Subhasmita Panda, Basanta Pravas Sahu, Soumya Ranjan Mahapatra, Jyotirmayee Dey, Namrata Misra, Rachita Sarangi

**Author notes:** Corresponding Author: Rachita Sarangi.

## Abstract

In addition to Coxsackie virus (CV), another pathogen that causes Hand-foot and mouth disease (HFMD), Enterovirus 71 (EV 71) is currently regarded as an increasing neurotropic virus in Asia and can cause severe complications in paediatric patients with blister like sores or rashes on the hand, feet and mouth. Not withstanding the significant burden of the disease, few treatments are currently available, and there is no authorised vaccine available for the disease prevention. Several vaccinations based on attenuated and inactivated vaccines have previously been identified, however they become worthless over time owing to changes in the viral genome. As a result, the goal of the study is to create an immunoinformatics and reverse vaccinology pipeline for predicting a multi epitope vaccine. A novel vaccine construct using B-cell derived T-cell epitopes from the virulent polyprotein and found the induction of possible immune response, in order to boost the immune system, aBeta-defensin 1 preproprotein adjuvant with EAAAK linker was added at the N-terminal end of the vaccine sequence. The immunogenicity of the designed, refined, and verified prospective 3D- structure of multi-epitope vaccine was found to be quite high with non-allergen, and antigenic property. The vaccine candidates bound to the TLR-3 in a molecular docking analysis and the efficacy of the potential vaccine to generate a strong immune response was assessed by means of an in silico immunological simulation. Computational analysis has shown that the proposed multi epitope vaccine possibility safe for use in humans and elicit an immune response, making it a promising tool against HFMD viral genome.

## Introduction

Rapid in silico informatics-based technique vaccinomics (Poland GA et al., 2009) has acquired significant attention with the recent backthrough in the sequencing of various pathogen genomes and protein sequence databases, as it integrates immunogenetics and immunogenomics with bioinformatics for the production of vaccines (Flower DR et al., 2008). The “vaccinomics” approach has already used effectively in the combat against disorders like multiple sclerosis (Bourdette DN et al., 2005) malaria (López JA et al., 2001), and tumors (Knutson KL et al., 2001). These approaches to vaccine development, however, tend to function through the discovery of human leukocyte antigen (HLA) ligands and T cell epitopes (Petrovsky N et al., 2002). Most investigations on the epidemiology, pathophysiology, and vaccine development for HFMD focus on the EV71 because it is now the prevalent genotype in severe (approximately 80%) and/or fatal (nearly about 93%) laboratory confirmed cases of HFMD in China (Xing W et al., 2014), studies on the epidemiology, pathophysiology, and vaccine development of HFMD have mostly centered on EV71. Serotypes A-F were attributed to EV71 based on the phylogeny of VP1. While serotype B and C are found all around the world, serotype D, E, and F are only found in India and Africa (Cox JA et al., 2017). The circulating EV71 strains differ from each other since they regulated by geographical location and the frequency of outbreak. The evolution of EV71 strains makes it difficult to develop a vaccine that is effective against all known strains and subtypes (Lei X et al., 2015). When constructing an EV71 vaccine candidate, it is crucial to choose a strain that can protect against the vast majority of different EV71 genotypes and sub-genotypes that are currently available (Bello AM et al., 2022).

Some of the most common causes of HFMD are strains of the EV71 family. Young children are particularly vulnerable to the neurological complications and even death that can results from these strains. There is currently no vaccine available to prevent severe HFMD that has been authorised by FDA. As HFMD has progressed into a potentially fatal epidemic, devasting the lives of young children in cyclical epidemics across the Asia-pacific, is a pressing need for a vaccine against EV71 to be developed (Yi EJ et al., 2017). Numerous MHC-restricted epitopes are used to create multi-epitope vaccination, which are then recognised by T-cell receptors (TCR) of multiple clones originating from different T-cell subsets, cytotoxic T lymphocyte (CTL), Th and B-cell epitopes inducing strong cellular and humoral immune responses simultaneously produce sustained immune responses and lower the risk of aberrant immunological reactions or adverse consequences caused by noxious substances (Lu IN et al., 2017; He R et al., 2018). In order to create a multi epitope vaccine that may induce a protective humoral and cellular immunological response against EV71, this study employs an immune-informatics driven screening strategy using EV proteome data.

## Materials and method

### 1. Selection of EV strain and extraction of protein sequences

The whole genome sequence of enterovirus A71 strain along with whole proteome, the NCBI database was searched for all encoded proteins(https://www.ncbi.nlm.nih.gov/protein). The polyprotein that plays crucial role in viral pathogenesis were chosen for vaccine candidate development, and their FASTA- formatted sequences were obtained from the UniProt database(http://www.uniprot.org/).

### 2. Forecasting of CTL epitopes

NetCTL 1.2 server (http://www.cbs.dtu.dk/services/NetCTL/) (Larsen et al.,2007),were used to screen the antigenic CTL epitope.For the prediction of CTL epitopes, the thresholds are 0.05, 0.15, and 0.75, respectively, for MHC-I binding affinity, artificial neural network (ANN) predicted proteosomal C terminal cleavage, and transporter associated transport efficiency. All the strong binding epitopes with IC50 value >500 was selected from the IEDB MHC-I binding resource.

### 3. Forecasting of HTL epitopes

T cell activation may be caused by the need of T cell receptors to epitopes complexed with the MHC class II molecule; Protein sequences were submitted to the NetMHCII Pan 3.1 server (http://www.cbs.dtu.dk/services/NetMHCIIpan-3.1/) and confirmed from IEDB research resource (http://tools.iedb.org/bcell/)in order to anticipate MHC class II restricted HTL epitopes. In order to determine the MHC-II epitope and allele binding affinity, a cut off value of 0.5% for strong binding peptides and subject to 2% value for weak binding peptides are used (Reynisson et al., 2020). Here, the strongest contender for the epitope was one that could bind to the most HLA-DR, DP, and DQ molecules. It was done using IEDB default parameter and can cover 99 % of the minimal allele in the HLA allele (Sette et al., 2008). HLA epitopes were predicted using a reference set consisting of 7 allele, and the species/locus was determined to be Human/HLA-DR. Additionally, the epitopes’ 15-mer length was obtained and classified based on percentile value.By percentile and IC50, the outcomes were ranked. High affinity is indicated by IC50 ≤ 50 nm, whereas intermediate and low affinities are indicated by IC50 ≤ 500 nm and IC50 ≤ 5000 nm, respectively.

### 4. Linear B-cell epitopes prediction

B cell epitopes are crucial for triggering a humoral immune response which in turns activate B cells to produce antibodies and plays a vital role in the development of vaccine. The B-cell epitopes can be classified as either linear or continuous and conformational or discontinuous depending on their structural configuration.The linear B-cell epitopes were predicted by ABCPred2.0 (https://webs.iiitd.edu.in/raghava/abcpred/ABC_submission.html) (Saha S et al., 2007). The residues are expected to be an epitope are those with score greater than the threshold value 0.5. Based on the solvent accessibility and flexibility the discontinuous B-cell epitopes were predicted by ElliPro an online server (http://tools.iedb.org/ellipro/) with the default parameters (Ponomarenko et al., 2008).

### 5. Identification of protective epitopes

Instead of relying on one tool, multiple tools can be used to predict epitopes with a higher level of accuracy. After comparing the identified B-cell, CTL, and HTL epitopes, we were able to choose the best segment with high overlap that gives epitopes the ability to elicit humoral and cellular immune response. The selected epitopes may cause a specific immune response as each epitope is identified by a number of servers. To obtain the score of the best conjugate IEDB server (http://tools.immuneepitope.org/mhcii/) was used. Antigenicity and allergenicity is a vital factor in the construction of a vaccine as a good vaccine candidate should be able to elicit an immune response without triggering allergic responses. AllerTop v2.0 (http://www.ddgpharmfac.net/AllerTOP/) (Dimitrov et al., 2014) and ANTIGENpro (http://scratch.proteomics.ics.uci.edu/) (Cheng et al., 2005) measured the allergenic propertied of the construct whereas the antigenicity of the vaccine was predicted using the VaxiJen online server (http://www.ddgpharmfac.net/vaxijen/VaxiJen/VaxiJen.html) (Doytchinova et al., 2007). It develops the K closest neighbours (KNN), auto- and cross-variance (ACC) transformation and amino acid E- descriptor machine learning algorithms for allergen categorization by investigating the physicochemical properties. The accuracy of this method is reported to be 85.3% during fivefold cross-validation (Dimitrov et al., 2014). Additionally, the SOLpro online tool (http://scratch.proteomics.ics.uci.edu/) was used to estimate the solubility when the protein construct was overexpressed in E. coli with a prediction accuracy of 74% and a 10-fold cross-validation approach (Magnan et al., 2009). Finally, the ToxinPred server was used to test all of the epitopes for toxicity(https://webs.iiitd.edu.in/raghava/toxinpred/algo.php) (Gupta et al., 2013).All the epitopes selected are nontoxic, antigenic and nonallergenic and should be induce the immune receptors. The percentile rank score is inversely related to the epitope binding affinity, i.e. the lower the percentile ranks, higher the binding affinity.

### 6. Interferon gamma (IFN-γ) epitope prediction

The IFN- inducing epitope server (http://crdd.osdd.net/raghava/ifnepitope/) was used to determine if the anticipated HTL epitopes would be able to trigger Th1 type immune response followed by the generation of interferon gamma by MHC-I activated CD4 + T helper cells.

### 7. Conservancy analysis of epitopes

In order to check the conservancy level of viral serotypes Epitope conservancy analysis service from IEDB (http://tools.iedb.org/conservancy/) was used (Bui et al., 2007) which assess the degree of epitope dissemination in the homologous protein set, that investigates the conservancy of chosen B cell, CTL, and HTL epitopes. The default values were utilized for the other analysis while the analysis type and sequence identity threshold were specified to “linear” and “100%”. To set the conservation tool to use as homologous protein, all Coxsackie family polyproteins were retrieved from the NCBI database. A vaccine would preferably have conserved epitopes that can elicit particular B-cell, and T-cell (CD4 and CD8) responses. In this study, we examined the conservancy of the selected epitopes of the polyprotein across different strains of enterovirus family.

### 8. Population coverage analysis

A vaccine candidate should provide a broad range of protection against different populations all around the world. Extreme polymorphism of MHC molecules (almost 6000) results in diverse pools or frequencies in people from various races and nationalities. Therefore, using a variety of peptides with various HLA binding abilities can enhance the population covered globally (Khan MT et al., 2021). The population coverage of the predicted T-cell epitopes was accessed by using IEDB population coverage tool which is crucial to develop the global vaccine candidate including all the specifications set to default.

### 9. Construction of muti-epitope vaccine candidate sequence

The stimulation of adaptive immunity is heavily reliant on T cell activation. The efficacy of the vaccine is enhanced when one adjuvant was added at the N-terminal end i.e., Beta-defensin 1 preproprotein. EAAAK, KK, AAY and GPGPG act as a linker to connect the B cell epitope, CTL and HTL respectively as well as a 6X histidine tag is added to the C-terminal end. Beta defensins are the antimicrobial peptides that thought to have an impact in epithelial surface susceptibility to microbial infection inducing activation and degranulation of mast cells by enhancing the protective immune response (Bensch et al., 1995).

### 10. Physicochemical properties and Secondary structure analysis

The ExPASy ProtParam online web tool (http://web.expasy.org/protparam/) was used to assess the physicochemical parameters of the final vaccine construct, including amino acid composition, theoretical PI, molecular weight, instability index, half-life (invivo and in vitro), aliphatic index and grade average of hydropathicity (GRAVY) (Gasteiger et al., 2005). The secondary structure of multi epitope vaccine construct was generated by online tool PSIPRED(http://bioinf.cs.ucl.ac.uk/psipred/) which also predict the transmembrane topology, transmembrane helix, fold and domain recognition (McGuffin et al., 2000). The antigenicity, allergenicity and toxicity of the construct protein was calculated using Vaxijen, AllerTop, and ToxinPred respectively.

### 11. Homology model and 3D structure prediction

The tertiary structure or three-dimensional (3D) model was modelled using Robetta server(http://robetta.bakerlab.org). The template structure for the specified amino acid sequence is determined using PSI-BLAST, BLAST, FFAS03, Robetta uses a comparative modelling approach to create the structure. If no template is detected, de novo Robetta fragment insertion method is used.

### 12. Refinement of tertiary structure and validation

The modelled structure was refined by GalaxyRefine server (http://galaxy.seoklab.org/cgi-bin/submit.cgi?type=REFINE). The server uses molecular dynamic simulation to achieve structural relaxation overall while substituting high probability rotamers for amino acids. Typically, the output given was consist of five refined models with various parameter scores, including GDT-HA, RMSD, Molecular Probability, clash score, poor rotamer and Rama preferred (Heo et al., 2013). To achieve the high quality of the projected structure, loop refinement and energy reduction were performed. SISS_MODEL web server was utilised to validate the quality of the improved structure. The primary purpose was to validate the protein 3D structure. It does this by estimating the entire quality score of the structure, which displays as a Z-score was less than the range of the characteristics for native proteins. Using the PROCHECK server (https://saves.mbi.ucla.edu/), a Ramachandran plot was generated that illustrates the energetically allowed and disallowed angles psi (ψ) and phi (φ) of amino acids (Laskowski et al., 1996).The server isolates the plots for glycine and proline residues and applies the PROCHECK concept to validate the protein structure, Ramachandran plot was used.

### 13. Disulfide bond engineering for vaccine stability

Whilst also changing cysteine residues in a highly mobile region of the protein, disulfide engineering can generate new disulfide bridges in the target protein. It significantly strengthens the geometric shape of the protein and are therefore essential for stability. Disulfide by Design v2.0 (http://cptweb.cpt.wayne.edu/DbD2/index.php) was used to engineer the disulfide bond, which searches for pairs of amino acid residues where cysteine can replace the original amino acid.

### 14. Discontinuous B-cell epitopes

Prediction of discontinuous B-cell epitopes of the vaccine design were performed using Ellipro tool on the IEDB server (Ponomarenkoet al. 2008). With a score of 0.8, 80% of the protein residues are located within the ellipsoids and 20% are located outside. Discontinuous epitopes are grouped together based on their distance R, denoted in Å, with a higher discontinuous epitope having a grater R value.

### 15. Molecular docking of final vaccine candidate with immune receptor

The interaction between an antigenic molecule and a specialized immune receptor is what triggers an adequate and efficient immune response. Molecular docking is a computer base method in which a ligand and a receptor interact to form a stable aggregate with a determined score indicating the degree of binding interaction (Shey et al., 2019). Toll like receptor 3 (TLR3) (PDB ID:1ziw) was downloaded from the Protein Data Bank as a receptor and the refined vaccine construct was used as the ligand. ClusPro (https://cluspro.bu.edu/login.php) online server was used to predict the interaction between the protein molecules. Water molecule and other small molecules were removed from TLR3 by using pyMOL.Results showing the lowest affinity score and prediction score is consider as the better score.

### 16. Molecular dynamic simulation

Then, MD simulation was performed on the highest-scoring complex found by protein-protein molecular docking, which involved discovering the optimum orientations for the vaccine candidate and the receptor proteins to interact with one another. GROMACS 2019 package and OPLS-e force field were used to do the MD simulation. The TIP3P water model was utilized in order to solvate the protein complex. Then, a neutral physiological salt concentration was achieved by adding sodium and chloride ions. Until a Fmax value of less than 10 kJ.mol was determined, the energy of the system was minimized using the steepest descent algorithm. The Linear Constraint Solver (LINCS) technique was used to impose constraints on all of the covalent bonds, ensuring that their lengths remained fixed throughout the simulation. The Particle Mesh Ewald (PME) approach was used to account for the long-range electrostatic interactions, and 0.9 nm was chosen as the threshold radius for the Coulomb and Van der Waals short-range interactions. In the next step, equilibrations were carried out for each system at 100 ps NVT (constant number of particles (N), volume (V), and temperature (T)) and 100 ps NPT (constant number of particles (N), pressure (P), and temperature (T). Then, GROMACS modules were used to do the necessary analysis, and xmgrace was used to generate the necessary charts and figures from the 50 ns of simulations run under periodic boundary conditions (PBC).

### 17. A5V codon adaptation, expression and purification

Codon optimization was used to increase expression of the recombinant protein. The degeneracy of genetic coding necessitates codon optimization because most amino acids can be represented by several codons. In order to assess the quantities of protein production in E. coli (strain k12), the JAVA codon adaptation tool (http://www.prodo ric.de/JCat) was used to acquire the codon adaptation index (CAI) values and GC content. A CAI of 1.0 is optimal, and anything above 0.8 is regarded excellent; the GC content is between 30% and 70%. Multi-epitope vaccine gene sequence optimization resulted in cloning into the E. coli plasmid vector pET28a (+). The sequence has been modified to include restriction sites at both the N and C termini, specifically for the enzymes EcoRI and BamHI. Snapgene software (https://www.snapg ene.com/free-trial/) was used to insert the final optimised sequence of the vaccine construct, including the restriction sites, into the plasmid vector pET28a (+).

### 18. Immune simulation

C- ImmSim (http://150.146.2.1/CIMMSIM/index.php) server was used to analyze the immune response of the vaccine construct, following cloning an in silico immune simulation was conducted. Three key elements of a functioning mammalian system- the lymph node, bone marrow, and thymus are simulated simultaneously. In real life, it is advised to wait for 4 weeks (28 days) between vaccination doses (Rapin et al., 2010). The interval between vaccine doses 1 and 2 should be at least 4 weeks. To simulate 1050 simulation steps, all primary parameter of C-ImmSim server were set to default and three injections containing 1000 vaccine proteins were given 4 weeks with time 1,8, and 168 for first, second and third injections respectively (each time step equals to 8 hours in real life, and time -step 1 is injection at time =0).

## Result

### 1. Analysis of retrieved protein sequence and profiling

In order to predict the B- and T-cell epitopes and design a multi epitope-based vaccine, the whole genome sequence of EV71 (under the accession number AAB39968.1) was obtained from the NCBI database, and the amino acid sequence for VP1, VP2, VP3, VP4, and polyprotein was obtained from the Uniprot database. The NCBI database revealed that human beta-defensin 1 (accession no. NP 005209.1) is an adjuvant that may stimulate an immune response against viruses. According to ExPASyProtparam’s analysis of the EV71 polyprotein’s physicochemical properties, the protein is stable with an instability index of 36.05, an aliphatic index of 83.15, and a GRAVY score of -0.198. VaxiJen 2.0 and AllerTop 2.0 were used to make the antigenicity and allergenicity predictions, respectively. With an antigenicity score of 0.5017, the protein was classified as a "Probable antigen," suggesting that it is likely not allergenic.

### 2. Cytotoxic T lymphocyte (CTL) epitope prediction

Using the NetCTL v2.0 server, we were able to predict nine-mer CTL epitopes from EV71, and then we narrowed it down to only the strongest binding epitopes by comparing it to the thousands of MHC-I epitopes provided by the IEDB server. Finally, we screened these potential epitopes using IC50 values below 200. **Table 1** lists the five antigenic and non-allergenic potential epitopes that were chosen among hundreds. Both the innate and adaptive immune systems rely heavily on interferon -γ (IFN-γ), a cytokine that has been shown to have antiviral, immunological regulatory, and anticancer effects. The IFN epitope server identified sequences capable of triggering IFN-γ.

**Table 1.** Prediction of CTL epitopes for multi epitope-based vaccine development.

| CTL Peptide seq. | Allele | Start-end | IC 50 | Antigenicity | Allergenicity | Immunogenicity |
| --- | --- | --- | --- | --- | --- | --- |
| HTDWANLVW | HLA-B*58:01 | 887-895 | 5.37 | 1.6136 | Non-allergen | 0.27686 |
| RSVPIGRAL | HLA-B*57:01 | 2169-2177 | 0.67 | 1.5002 | Non-allergen | 0.23028 |
| GSLEVTFMF | HLA-B*58:01 | 436-444 | 22.2 | 1.4707 | Non-allergen | 0.17009 |
| TINYTTINY | HLA-B*35:01 | 24-32 | 80.12 | 1.2485 | Non-allergen | 0.17706 |
| FADVRDLLW | HLA-B*58:01 | 996-1004 | 11.93 | 1.1216 | Non-allergen | 0.10418 |

### 3. Helper T lymphocyte (HTL) epitope prediction

The viral polyprotein was used to predict 29 non-allergenic, antigenic, and non-toxic HTL epitopes (15-mer). Out of them, only 9 were determined to have IFN-γ positive epitopes. Dual human allele epitopes with no overlap HLA-DRB1*04:01, HLA-DRB1*13:02, HLA- DRB1*08:02, HLA-DRB3*02:02, HLA DRB1*07:01, HLA-DRB1*01:01, HLA-DRB1*04:05, HLA-DRB1*11:01, and HLA-DRB1*09:01 were chosen as HTL epitopes for vaccine development because they have the IC50 values of the chosen epitopes were lower than 100. If the vaccine epitope has a low IC-50 value (often 100 is used as a cutoff), then it will be safer for use in humans at lower doses (Berrouet, C et al., 2022). **Table 2** demonstrates that all verified epitopes are non-allergenic, non-toxic antigens.

**Table 2.**
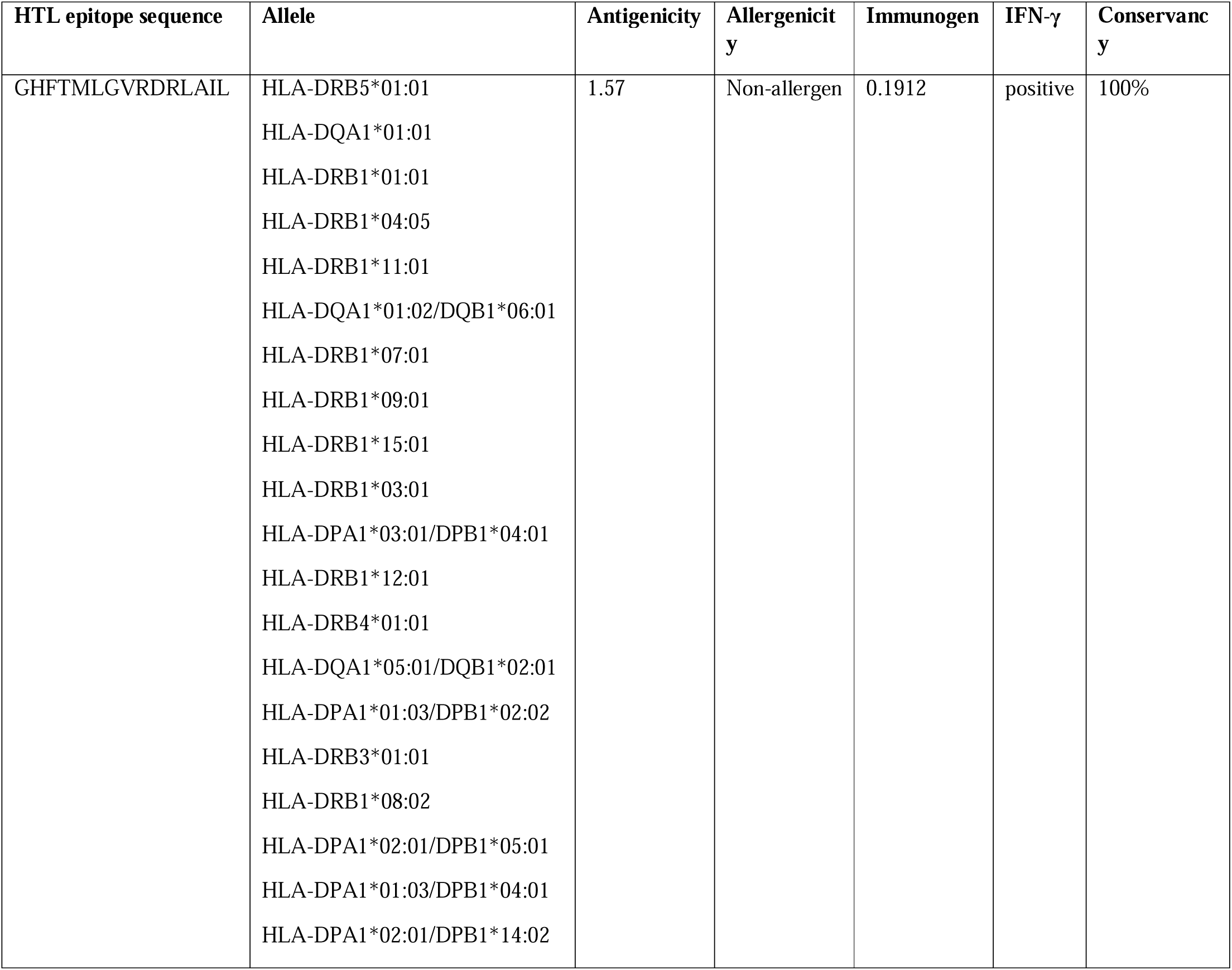
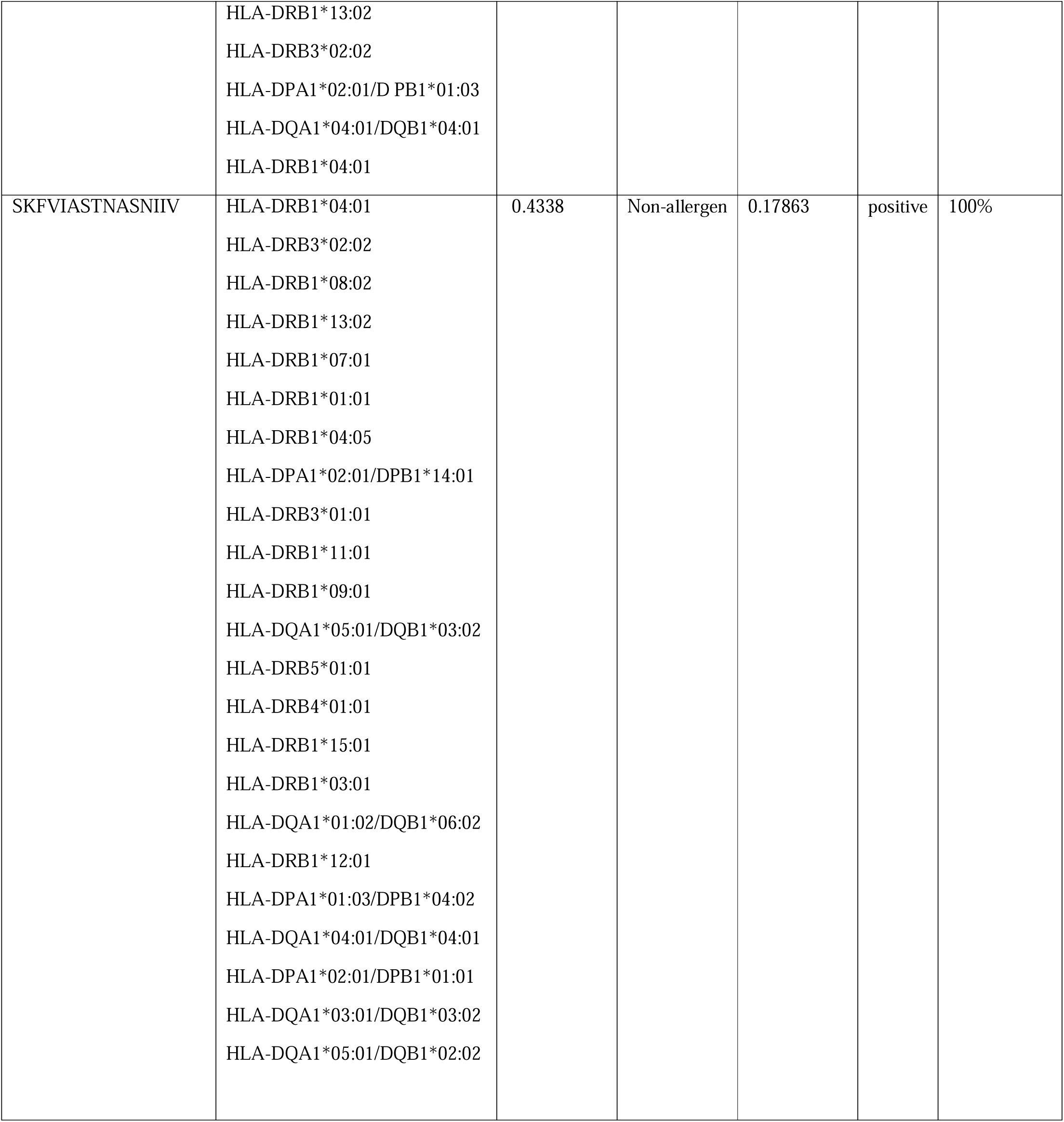
Prediction of HTL epitopes for multi epitope-based vaccine development.

**Table 3.** B-cell epitopes for multi epitope-based vaccine development.

| Peptide B cell epitope | Antigenicity | Allergenicity | Toxin pred |
| --- | --- | --- | --- |
| RIEPVCLIIR | 2.623 | Non-allergen | Non-Toxin |
| FKSKHRIEPV | 2.092 | Non-allergen | Non-Toxin |

**Table 4.**
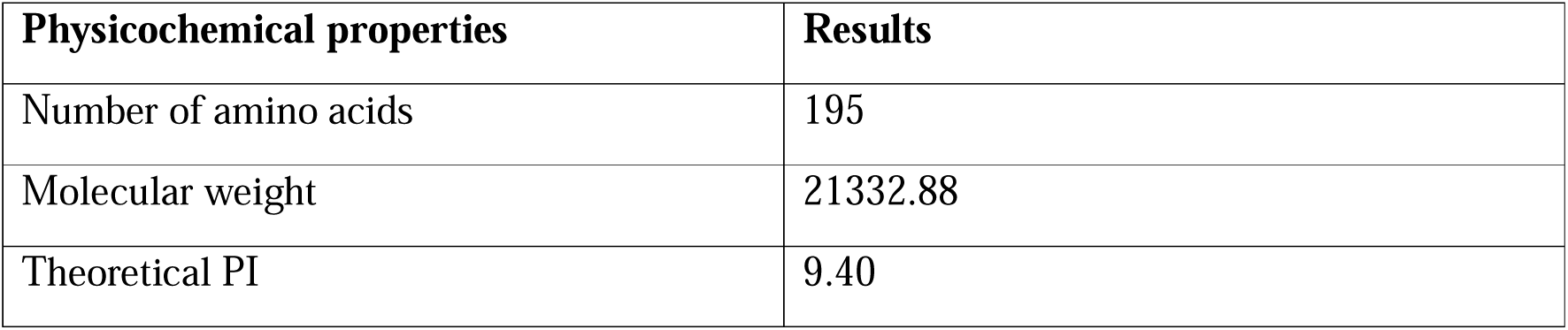
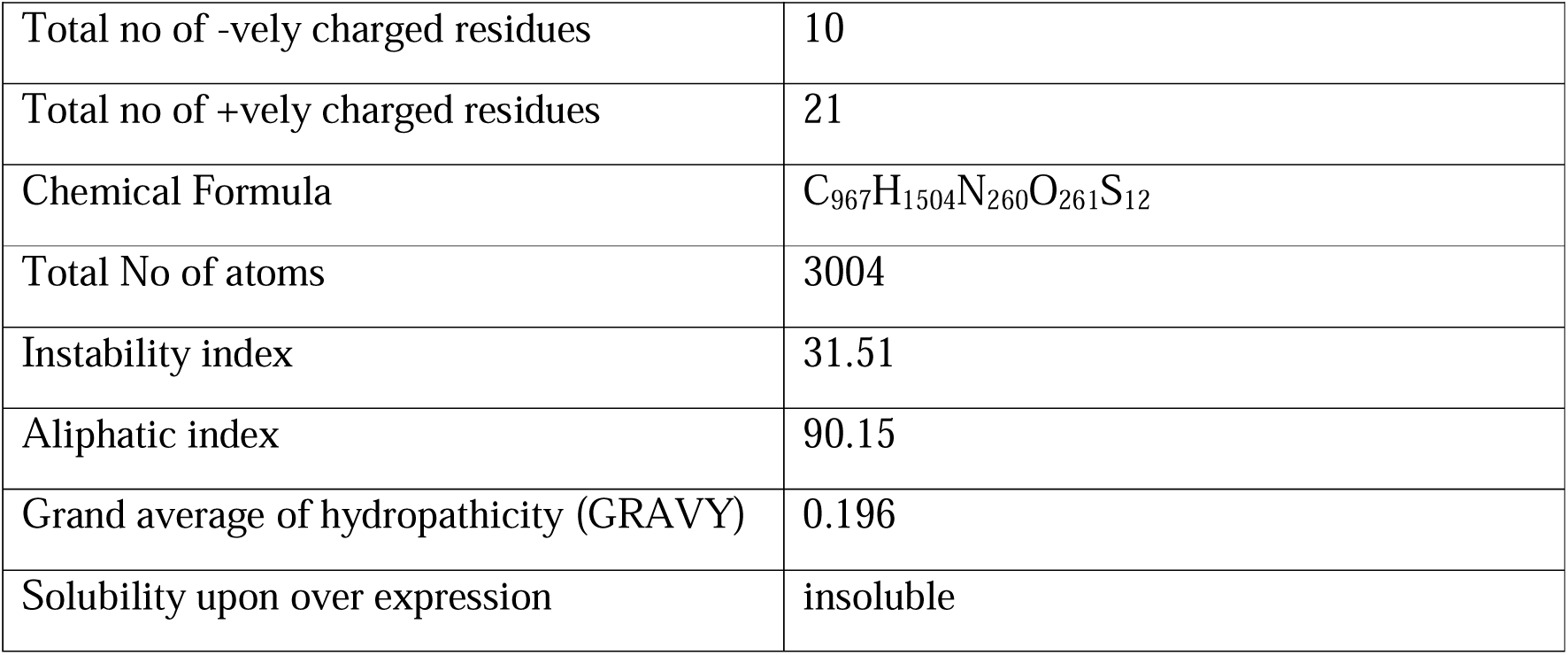
Physicochemical properties of vaccine construct protein

### 4. B-cell epitope prediction

Using IEDB’s linear epitope prediction tool v2.0 and artificial neural networking trained on fixed-length patterns of B-cell epitopes and random non-epitopes, the ABCPred was able to accurately predict the B-cell epitope for the multi-epitope vaccination with an accuracy of 65.93%. The greatest and lowest antigenicity scores for the projected epitopes are 1.741 and 0.587, respectively, indicating that they are all antigenic, non-allergenic, and non-toxic (Table-3).

### 5. Designing of a multi-epitope vaccine construct

Initiating adaptive immunity is highly dependent on T-cell regulation. Two B-cell epitopes, five CTL epitopes, and two HTL epitopes were nominated in order to design a novel vaccine that met the criteria of binding affinity, antigenicity, non-toxicity, and non-allergenicity. To increase the vaccine’s efficacy, human -defensin 1 was added to the N-terminus as an adjuvant. The construct’s stability was ensured by the addition of four linkers. The linker EAAAK was used to join the adjuvant to the B-cell epitopes. Nevertheless, linkers KK, AAY, and GPGPG were used to connect the B-cell, CTL, and HTL epitopes, in that order **(Fig-2)**. The vaccine design was projected to be non-allergenic based on an antigenicity score of 0.828 at a 0.4 threshold. The whole vaccine constructions have 195 amino acid residues, with the formula C967H1504N260O261S12. A 21kDa molecular weight and an isoelectric point of 9.40 characterize the synthetic vaccination. The half-lives in humans (at 30 hours), yeast (10 hours), and E. coli (10 hours). With an instability score of 31.51, the protein is considered to be stable. If the Gravy value is positive, it means that the polar character of the construct vaccine has been shown. The aliphatic index and Gravy value are 90.15 and 0.196, respectively **(Table-4)**. Overexpression of the construct resulted in insolubility, as anticipated by SOLpro (solubility was 0.673).

**Figure 1.**
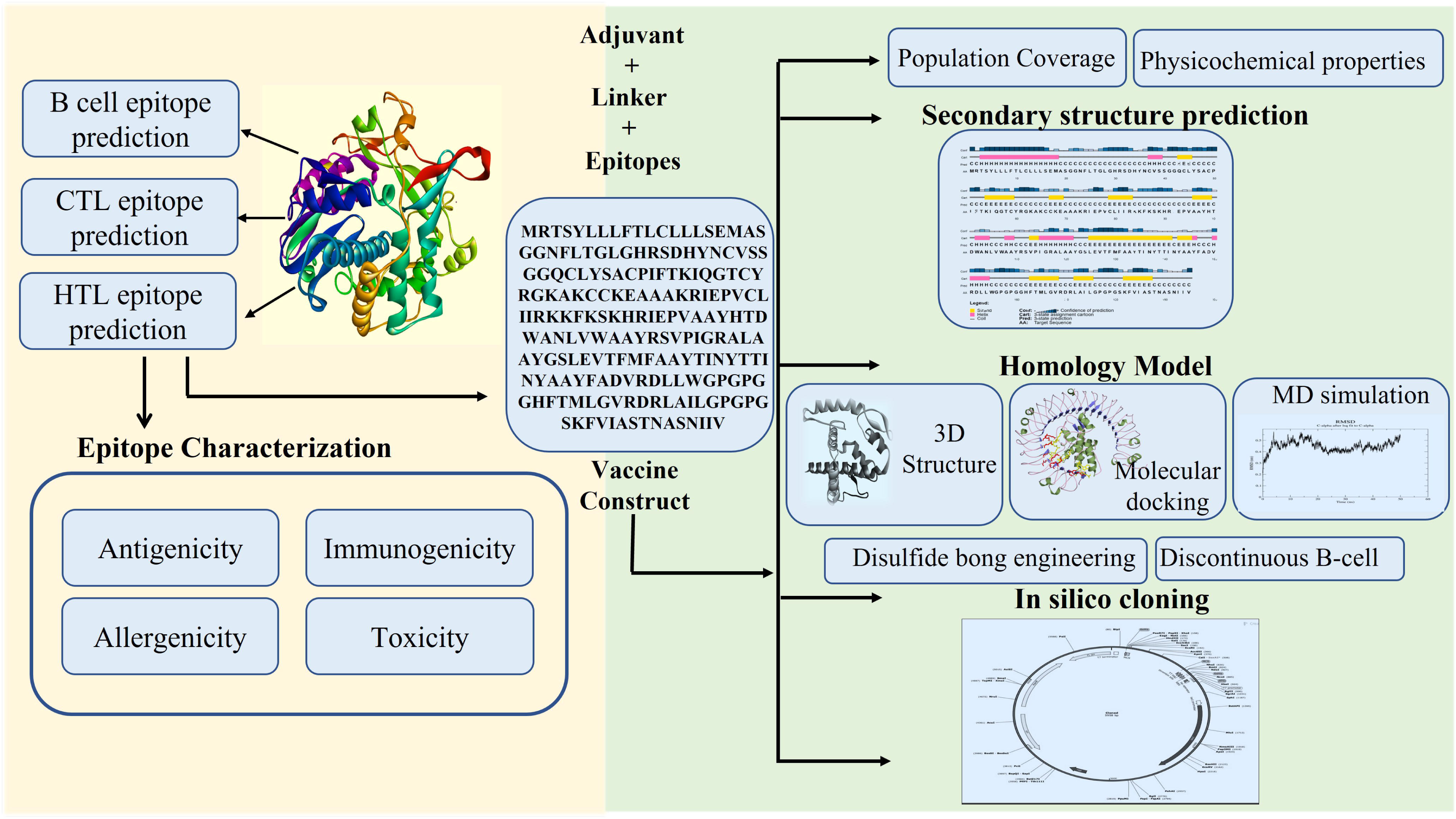
Flow chart showing the structural arrangement of designed vaccine.

**Figure 2.**
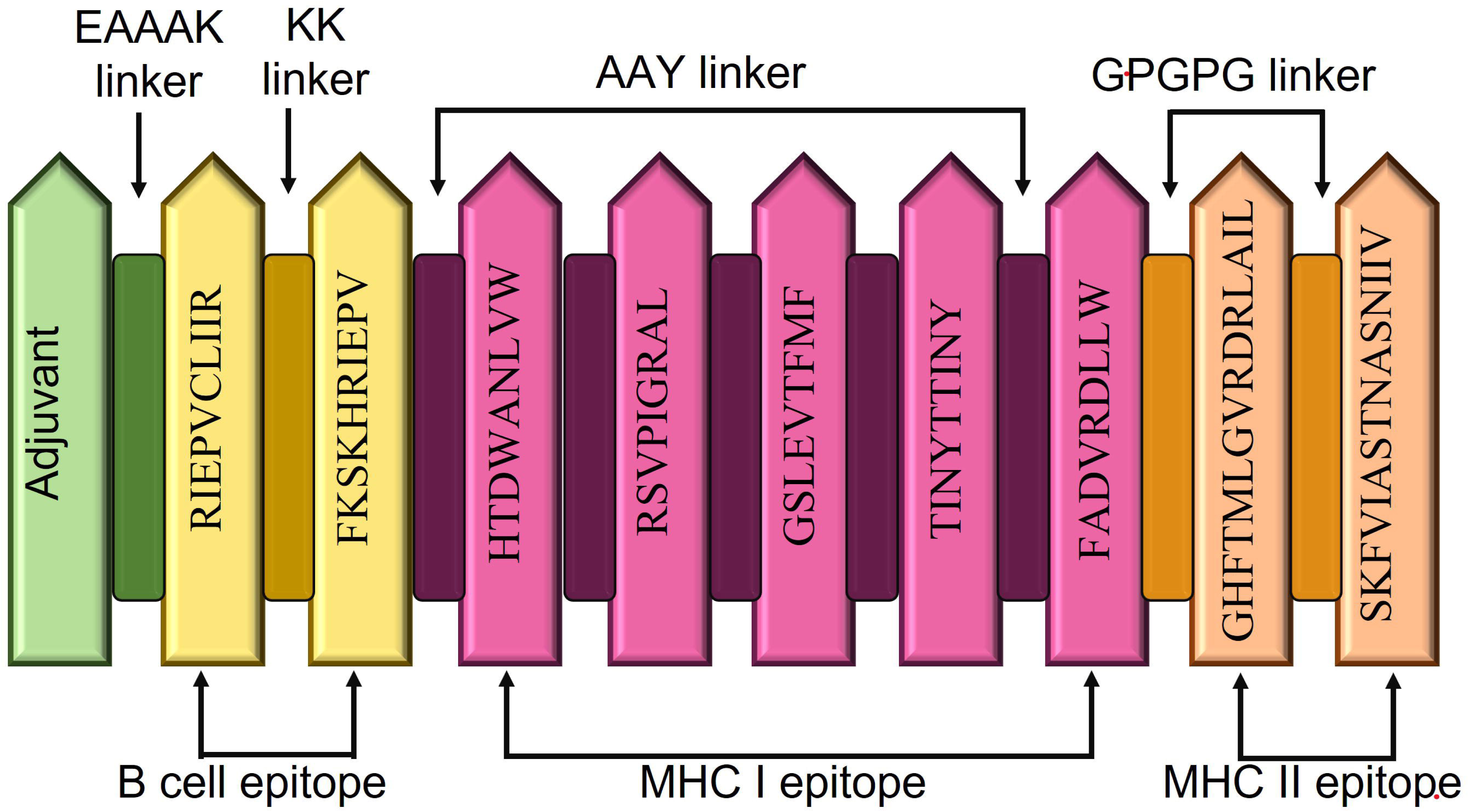
Schematic representation of the final vaccine construct. The vaccine construct contains 191 amino acids: Beta-defensin 1 adjuvant at the N-terminal region following B- cell, CTL and HTL epitopes with dissipated linkers EAAAK, KK, AAY and GPGPG respectively.

**Figure 3.**
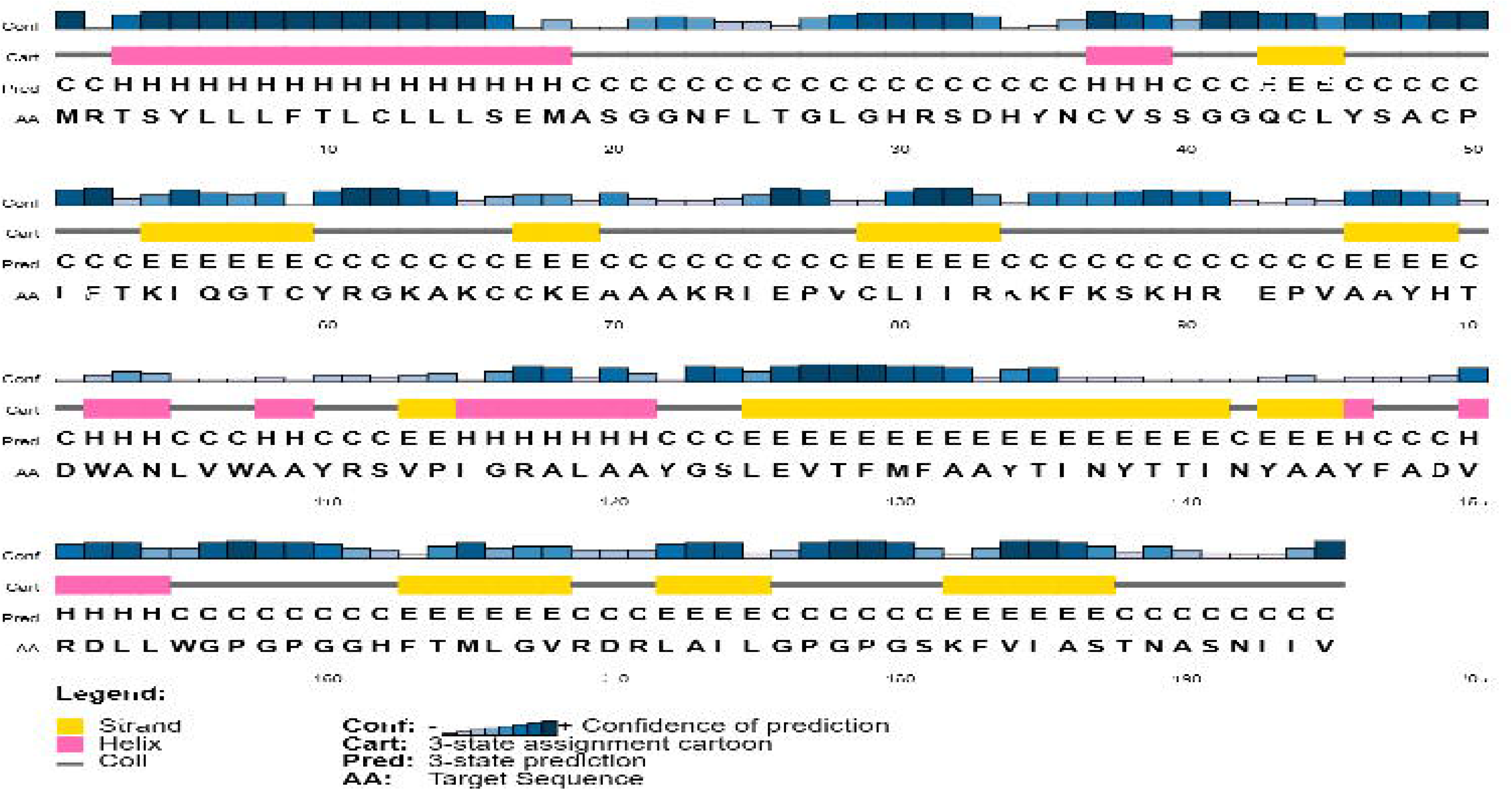
Secondary structure of the vaccine construct showing alpha helix, beta sheet and coil-coil region

### 6. Structure analysis, refinement and validation

Alpha helices accounted for 33.85% of the expected secondary structure, followed by extended strands (25.13%), and random coils (32.31%). Robetta, among the top five models, was used to create the major tertiary structure of the vaccine construct, with a higher C score indicating more model confidence. So, the 3D model was improved further by the GalaxyRefine server, yielding five final models for the vaccine build. Model 1 has the best refined structure according to a variety of criteria including a global distance test-high accuracy (GDTHA) of 0.978, a relative mean square deviation (RMSD) of 0.301, a Mol Probity of 2.167, a clash score of 14.6, a bad rotamer of 0.6, and a Rama favoured of 91.7%. As a result, model 1 was chosen for further studies. We ran the 3D structure via the PROCHECK and SWISS-Model server, where we got the Ramachandran plot and found that 91.7% of the residues were in the preferred area. Ramachandran plots show which dihedral angles (psi and phi) are permissible and which are not allowed from an energy perspective. In addition, ERRAT, verify 3D, and 84.10 percent of the time showed that the model driven after GalaxyRefine had better stereochemical quality, validating the quality evaluation.

### 7. Epitope Conservancy analysis

Using the whole genomes of isolates, we were able to determine the polyprotein amino acid sequences that are responsible for their infectiousness. The IEDB conservancy analysis tool was used to examine the conservation of the chosen B-cell, CTL, and HTL epitopes across all strains of the enterovirus family. More than 80% of the chosen epitopes were shared across the isolates from various nations, whereas 4 antigenic epitopes exhibited full 100% conservation.

### 8. Population coverage

Only those with the appropriate MHC molecule will have an immune response to a certain epitope. Increased coverage of the patient population targeted by epitope-based vaccinations or diagnostics may be achieved by selecting several peptides with various HLA binding specificities. Based on their worldwide distribution features, the identified T-cell epitopes are ideal for inclusion in the construction of the candidate protein vaccine, as shown by the allelic frequency statistics. Finding the MHC allele linked with the best epitope set for population coverage is important. The population coverage of the predicted epitopes, demonstrating that the chosen epitopes covered 95.6% of the global population in 109 nations across 16 regions. The percentage of the world’s population covered was observed to be above 90% in almost all nations.

### 9. Disulfide bond engineering

The analysis showed, for disulfide bond engineering a total of 124 pairs of residues were exploited among that only those pair of residues fulfilling all the criteria i.e., the energy and χ3 values must be within 2.2 and between -87 to +97° respectively will be accepted. As a result, only one pair of residues were fulfilling all the criteria, therefore one mutation was generated on the residue pair ALA48-ASP152 with a χ respectively. The disulfide bonds with blue and pink colors are shown in the created mutant (Fig.6).

**Figure 4.**
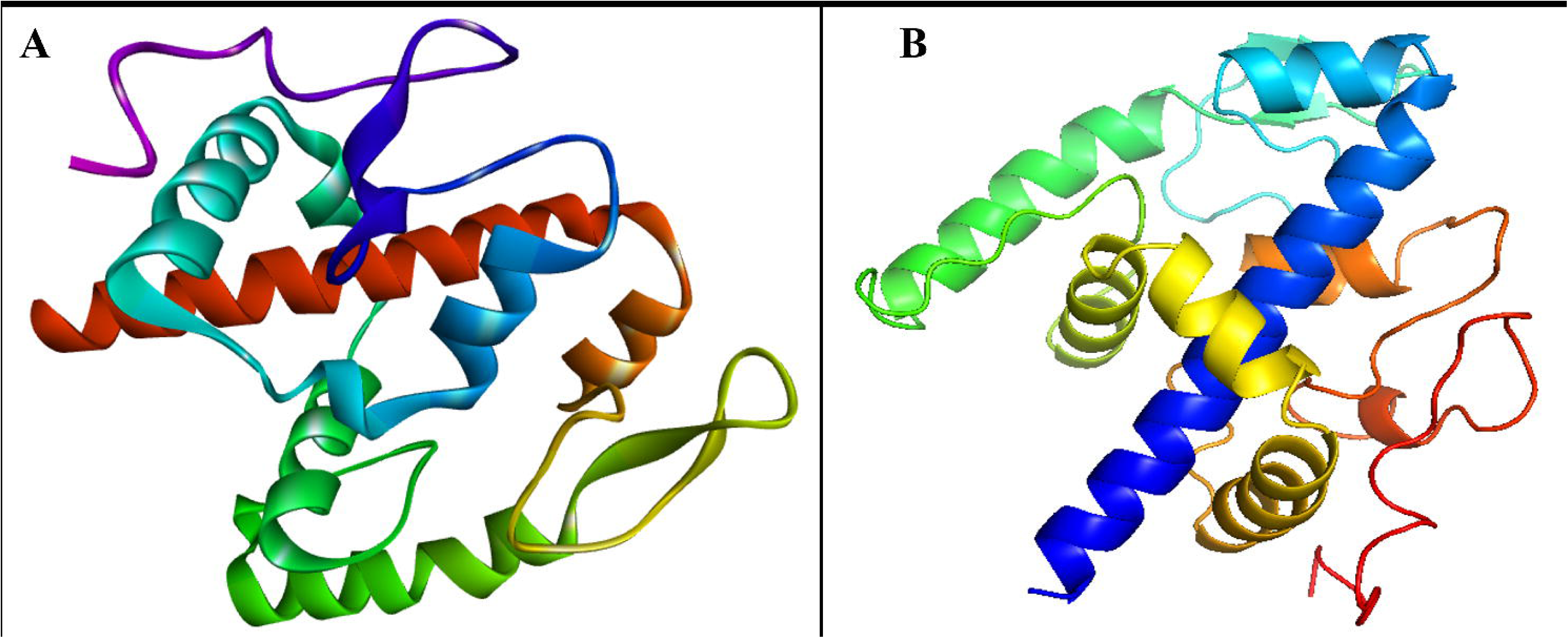
A. Homology modelling of the vaccine. **B.** Refinement structure of the final vaccine construct.

**Figure 5.**
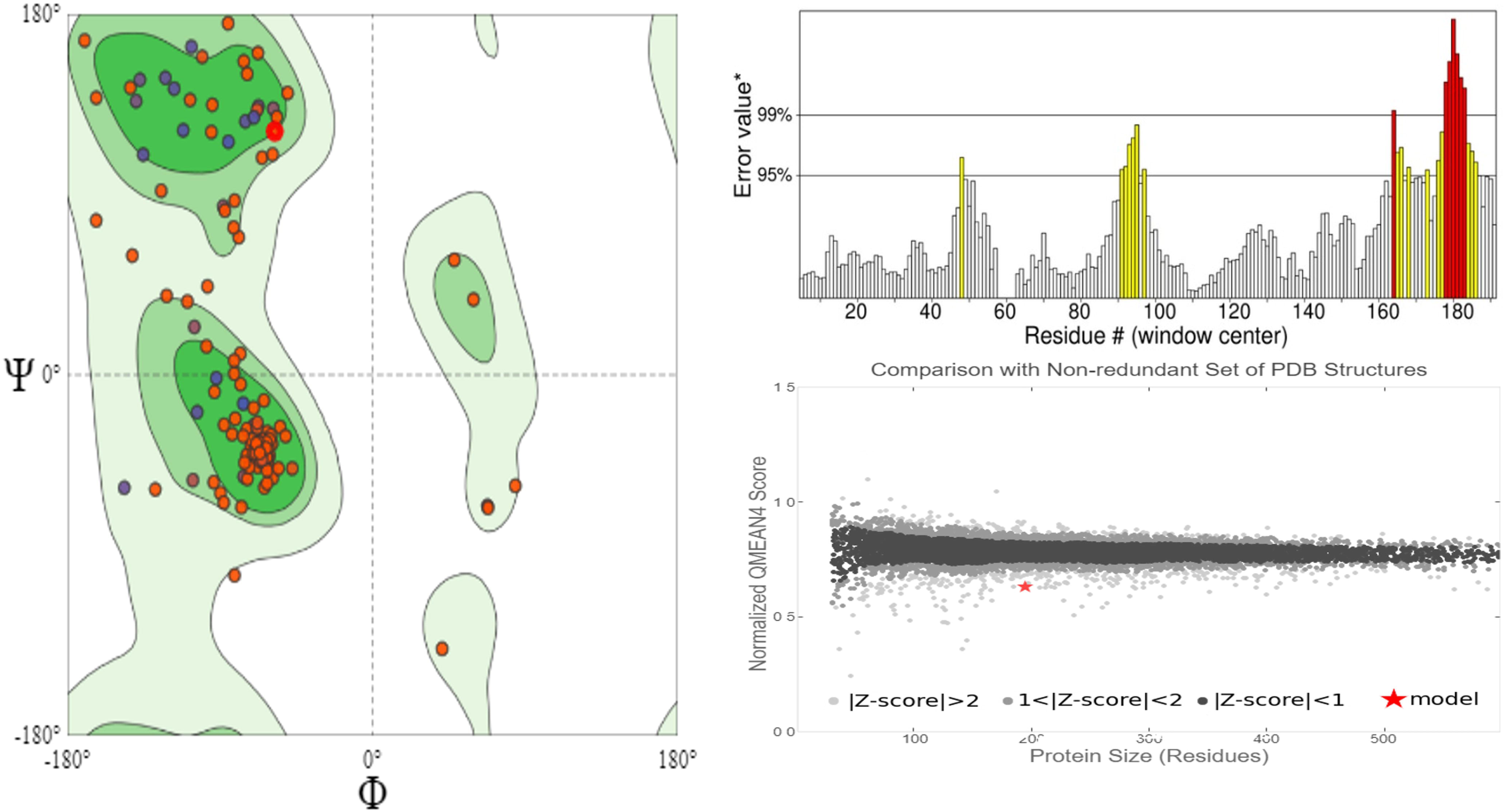
A. assessment and validation of final model by SAVES web server**. B.** Ramachandran plot analysis of refined structure showing amino acid residues in the favoured, allowed and outlier region.

**Figure 6.**
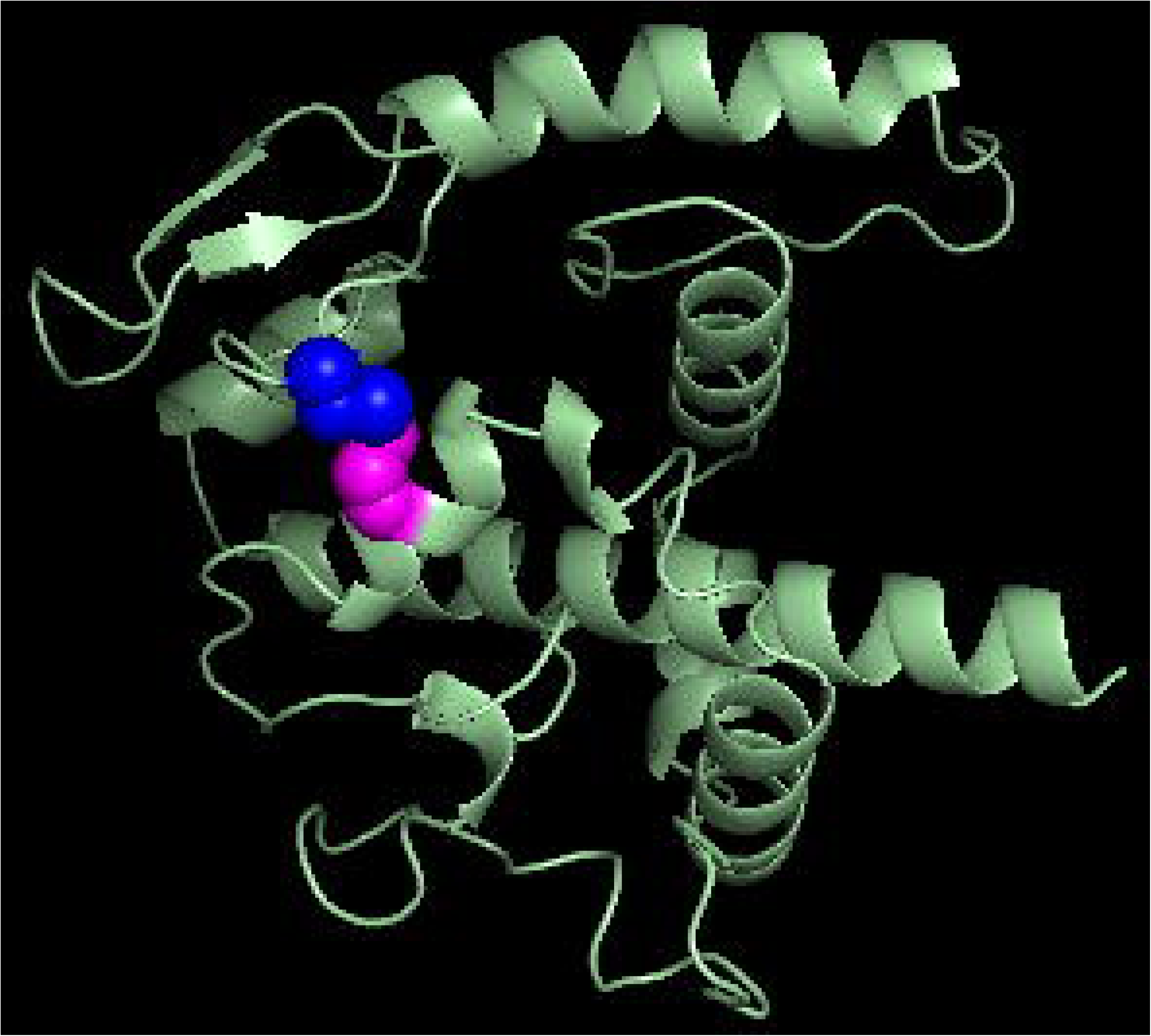
Disulfide bond engineering of the vaccine construct mutant model showing the disulfide bond formation.

**Figure 7.**
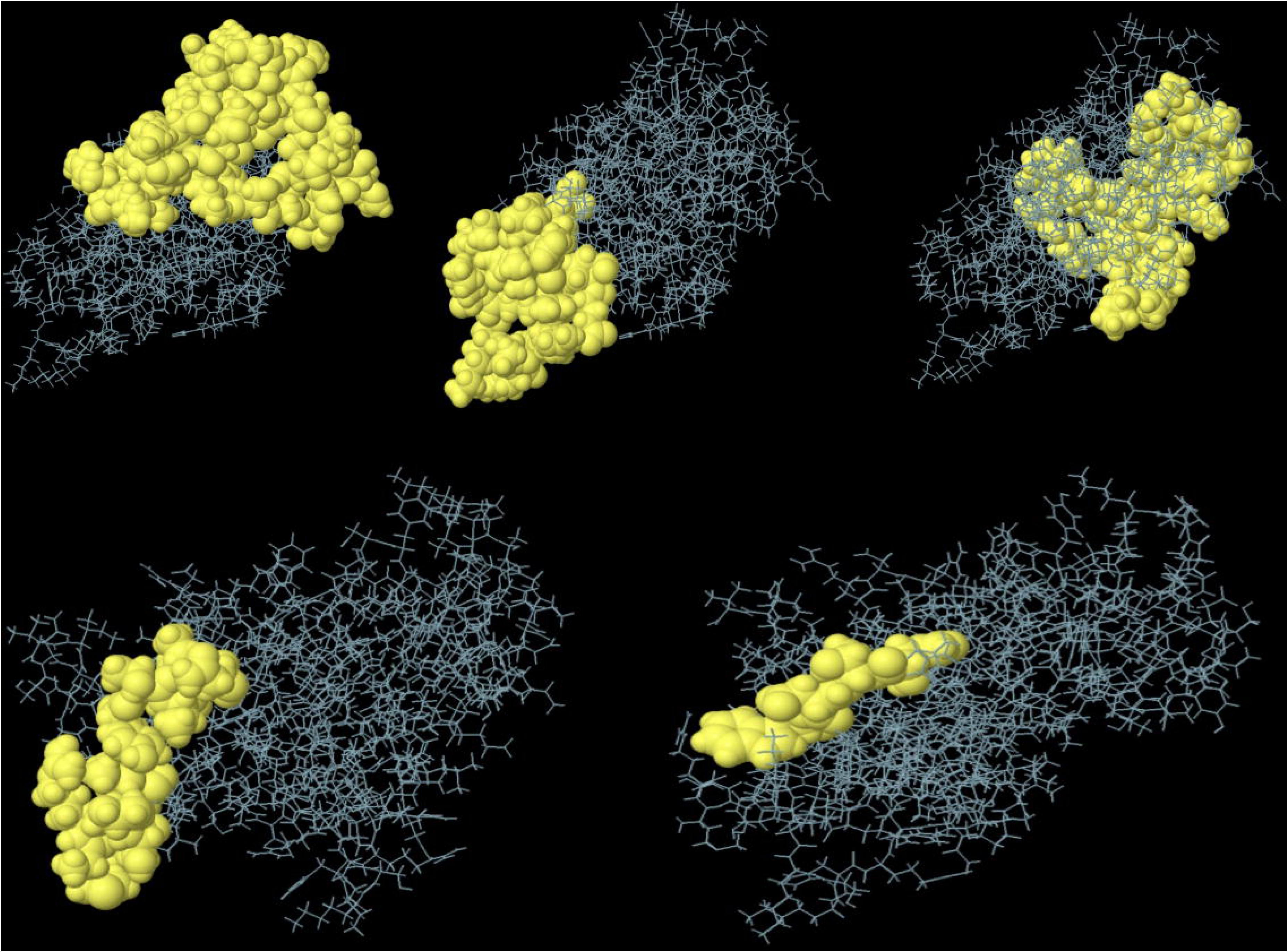
Predicted confirmational B-cell epitopes by Ellipro tool.

**Figure 8.**
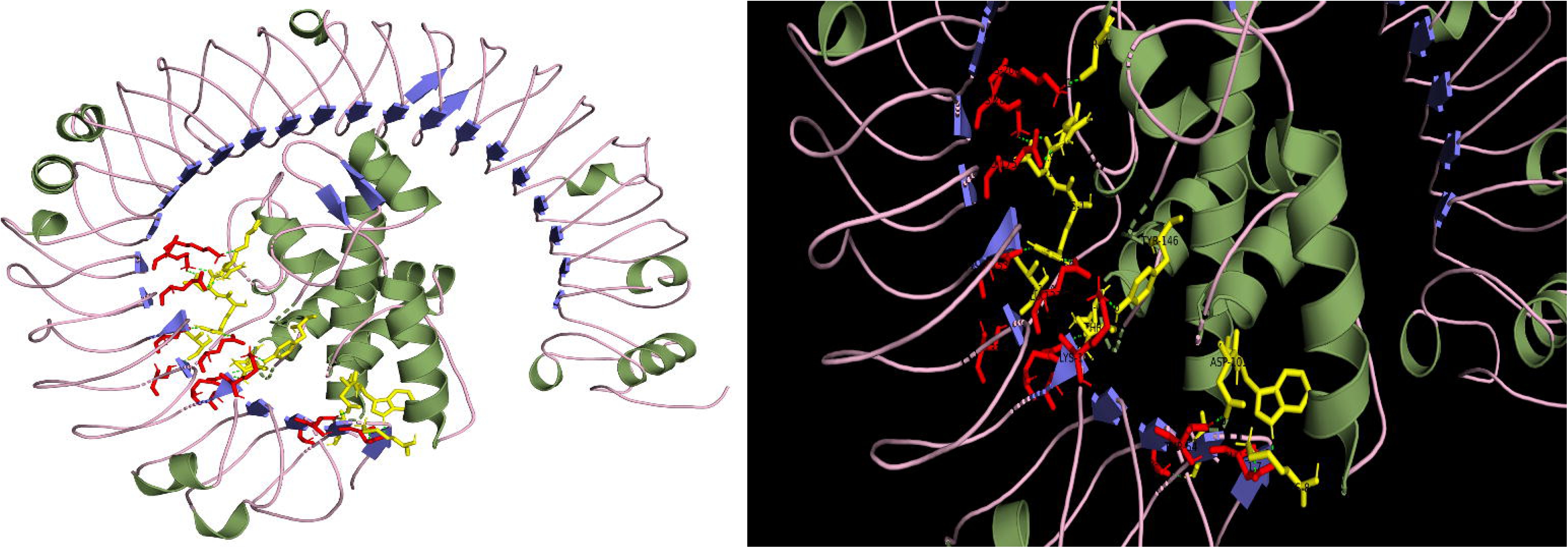
Molecular docking of final vaccine construct with TLR-3 complex and hydrogen bond interaction map.

**Figure 9.**
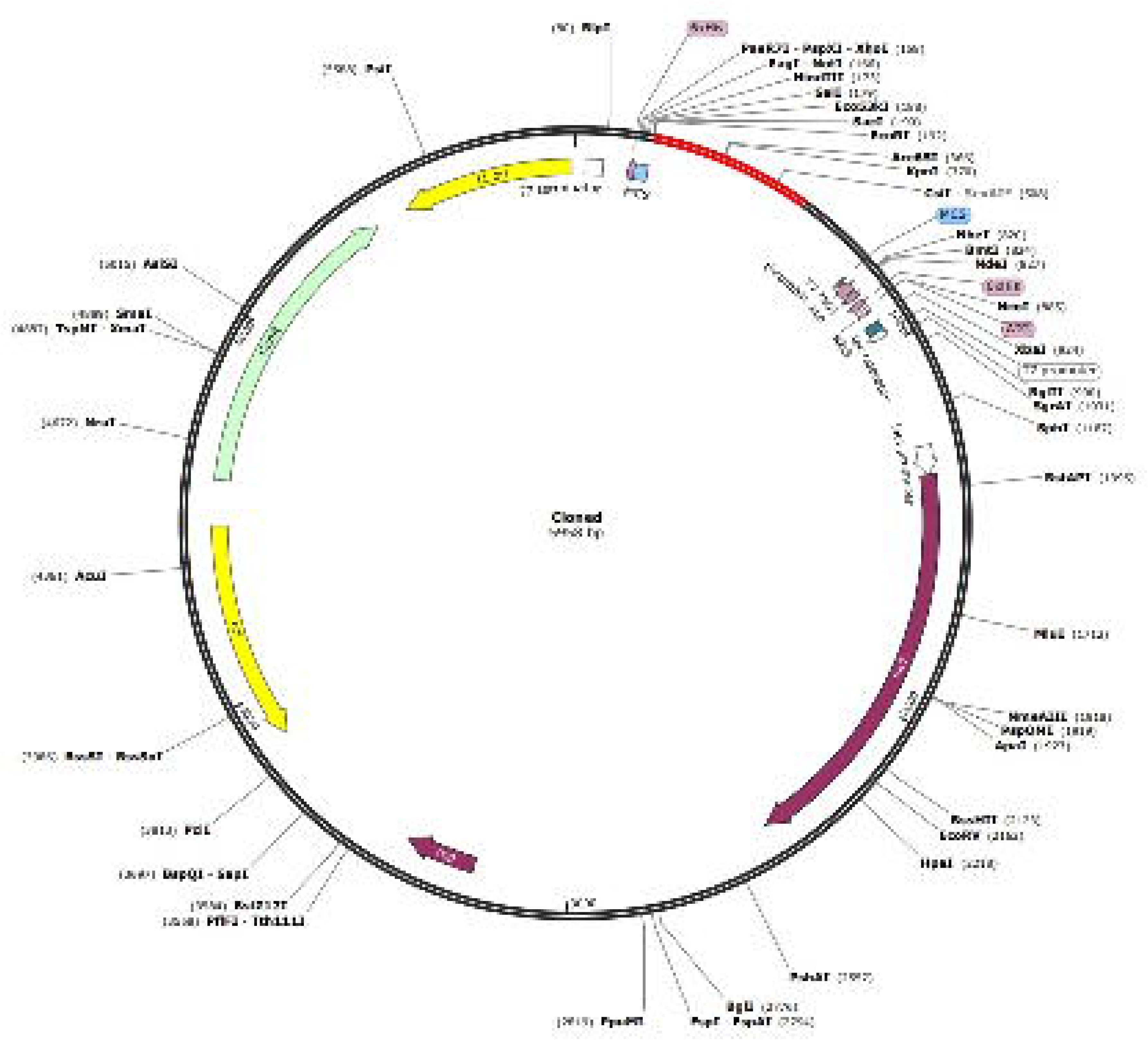
Molecular dynamic simulation showing Root mean square deviation and root mean square fluctuation analysis of protein backbone and side chain residues.

**Figure 10.**
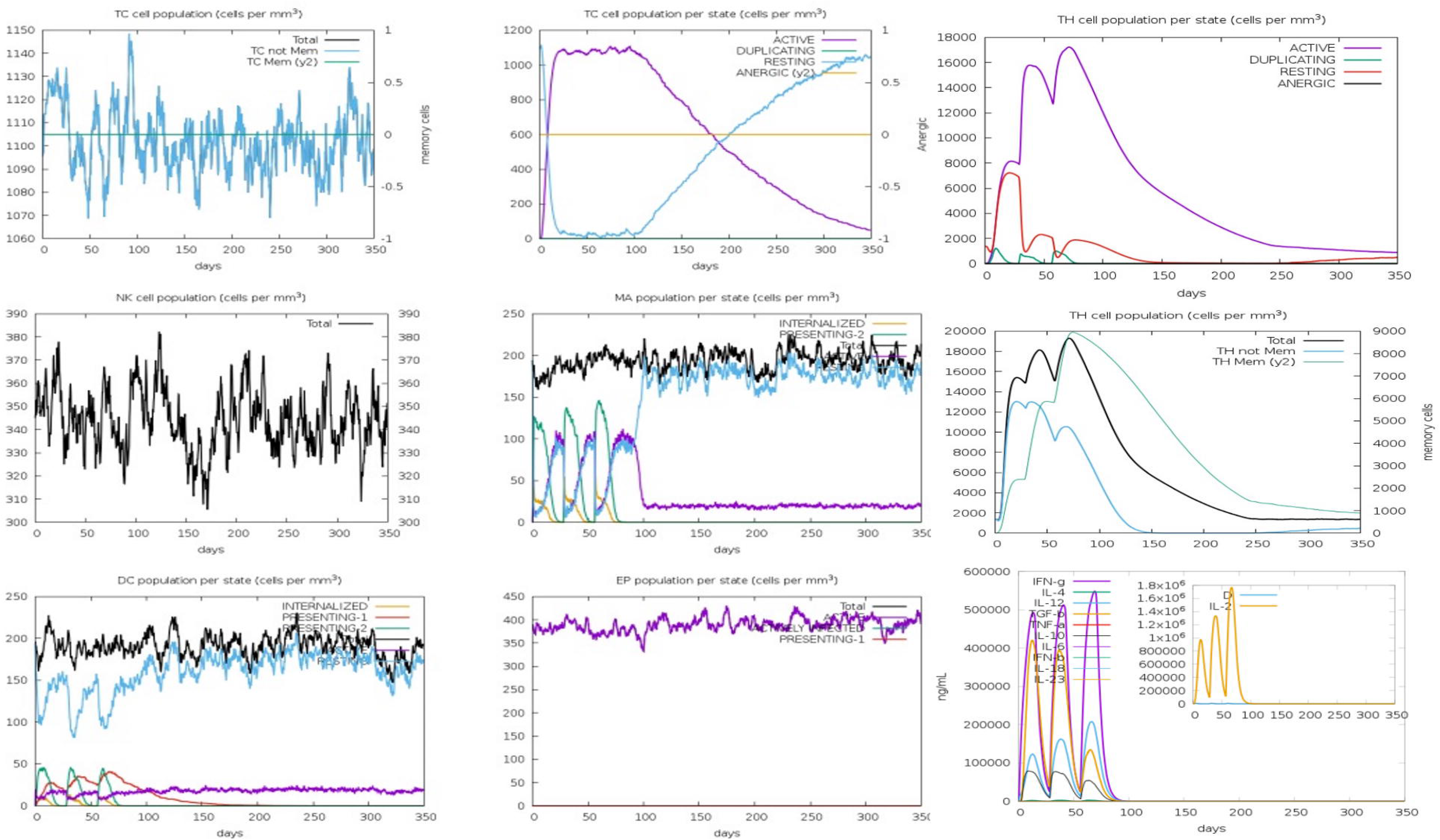
In silico cloning of the gene sequence in to the pET28 (+) expression vector. The red part showing the gene coding for multi epitope vaccine construct and the black region showing the vector backbone.

### 10. Conformational B-cell epitope prediction

Four discontinuous B-cell epitopes contain115 amino acid residues which values are ranging from 0.50-0.78 and the size ranging from 4-40 residues were estimated to be suited in 5 discontinuous epitopes (Fig-). From vaccine sequence length different discontinuous epitope residues were predicted such as 76-94, 98 (20 residues); between 26,27, 29-36, 39-42, 44, 46-50, 52-66, 68-70, 72-74 (40 residues); between 114-116, 122-127, 129-130, 134, 137, 156-161, 174-195 (40 residues), between 1-6, 138-142 (11 residues) and between 99-102 (4 residues) have been predicted. The individual scores were shown in **Table-5.**

### 11. Molecular docking

The immunological response to Ev-71 is bolstered by toll-like receptor 3. (Chen GP et al., 2021). So, using ClusPro, we docked the final vaccine construct to TLR-3 (PDB ID:1ziw), and the resulting complex had an energy score of -1183.2 when evaluated in Pymol, making it the best possible candidate for an effective vaccination. Many strong hydrogen bond (H-bond) connections with reduced distance between the interacting residues were seen between the vaccine construct and the immunological receptor, demonstrating high binding affinity. The residues of docked vaccine-TLR3 complex showing H-bond interactions with distance were documented in **Table-6** (**Fig- 8**).

### 12. Molecular dynamic simulation

The optimal molecular docking complex was used to drive the MD simulation. After this, a 50-ns MD simulation was performed on the complex, and the results were analyzed in terms of the root mean square deviation (RMSD), the root mean square fluctuation (RMSF), the number of hydrogen bonds (H-bonds), and the radius of gyration (Rg). The relative mean square deviation (RMSD) profile represents the changes in the protein backbone from the starting structural conformation to the final state. When taking into account all of the -atoms, the calculated average RMSD value for the receptor-vaccine combination was 0.4 nm (Figure-9). This suggests that positive interactions may lead to the formation of a stable vaccine-receptor complex. Using the RMSF, one can learn about the residue-by-residue dynamics of a protein relative to its initial state. Analysis of the RMSF of the protein atoms allowed us to learn about the conformational behavior of the ligand-receptor combination at the residue level. Estimated Mean Squared Fluorescence (RMSF) Value: 0.3 nm (Figure-9). Hydrogen bonding is essential for protein stability and for the recognition of ligands by proteins. The amount of hydrogen bonds formed in the vaccine and receptor complex during the MD simulations was also analyzed to provide insight into the selectivity of interactions and the efficacy of the vaccine. on average, 15-18 hydrogen bonds were formed in the vaccine-receptor complex (Figure). As calculated by molecular dynamics simulation, the vaccine-TLR-3 complex has an average radius of gyration (Rg) of 3.15 nm (Figure-9). This research demonstrated that after a productive contact between the vaccination protein and the TLR-3 receptor, the complex became more compact.

### 13. Codon optimization and in silico cloning

The vaccine design in E. coli strain k12 has its codon use optimized using the Java codon adaptation tool for maximum expression. Excellent expression of the vaccine candidate in E. coli host was predicted thanks to a CAI of 1.0 for the optimized nucleotide sequence and a GC content of 52.30 for the modified sequence. Cloning the vaccine construct will be easier with two restriction endonucleases, such as EcoRI and BamHI, placed at opposite ends of the DNA. The altered codon sequences were then inserted into the plasmid vector pET28a (+) with the aid of SnapGene software, yielding a recombinant plasmid sequence of 5958bp in length (Figure-10).

### 14. Immune simulation

Evaluation of immunogenicity occurs at several stages throughout vaccine candidate development. The real immunological response of the produced vaccination in the mammalian system is predicted by the C-ImmSim server. To conduct this investigation, the amino acid profile, epitopes, and immunological responses were mostly used. The initial immunological response was characterized by a high level of IgM. Secondary and tertiary immune responses are characterized by an increase in B-cells and by the production of large amounts of IgM, IgG1+IgG2, and IgM+IgG despite a reduction in antigen concentration (Shey et al. 2019). The development of immunological memory is marked by the appearance of new B-cell isotypes as well as B-memory cells. The numbers of TH and Tc cells have grown in tandem with the maturation of memory T cells. We observed an upsurge in IFN-g, TGF-b, IL-10, and IL-12 and a reduction in IL-4, IL-6, IL-18, IL-23, TNF-a, and IFN-b. An adequate amount of immunoglobulin is produced when the Simpson index (D) is low and there is high expression of dendritic cells (cell-mediated immunity) and IFN-g and IL2 (humoral immunity).

The smaller D value indicates the lower diversity of T cell clones (Figure. 11). The immune simulation study was conducted to understand the antigenicity of the vaccine and also to reveal the generation of adaptive immunity and immune interactions. In the immune simulation analysis, we proved that our proposed vaccine candidate can act as a prominent antigen because they have the ability to induce the production or generation of antibodies. From the overall analysis, i.e., based on the immunological responses; we can confirm that the human immune system recognizes the polyprotein from the first day onwards, and antibody production begins from the day ∼4th onwards. The analysis of cytokine and interleukin production is crucial for studying both innate and adaptive immunity; in comparison to other cytokines, IFN-g production is high from the 2nd day onwards, and it plays an important role in macrophage activation, as well as stimulating natural killer cells and neutrophils. As seen in the B-cell population (cells per mm3) analysis, our vaccine does not activate B isotypes IgG1 and IgG2, but they do activate B memory (y2) cells and B isotype IgM. Tc cell activity is very low, there is no Tc memory (y2) cell formation, or their activity is in a resting condition (Fig.11). During the formation of Ig, the formation of TH cells is also very important; from the 2nd day onwards, the TH memory (y2) cell population increased until it reached 9000 memory cells per mm3. However, the activation of TH cells was only observed from day 5th onwards (Fig.11). And the NK cell population is highly fluctuating in nature. Macrophage cells were stable throughout the time period. The vaccination also stimulated the interferon-γ, TGF-beta, interleukin-10, and interleukin-12 production (Fig.11D). There was an increase in the number of interferon-γ, TGF-beta, interleukin-12and interleukin-10 concentration after the first two vaccinations, however; their concentration decreased after the third vaccination in comparison to the first two. Overall, the C-IMMSIM simulations predicted that the multiepitope vaccine could activate the immune response.

**Figure 11.** In silico immune simulation by C-ImmSim using vaccine as an antigen **A.** Cytotoxic T-cell population per state **B.** Cytotoxic T-cell population **C.** Natural killer cell population **D.** Macrophage cells population per state **E.** Dendritic cells population per state **F.** Epithelial cells population per state **G**. Helper T-cell population per state **H.** Helper T- cell population **I.** Concentration of cytokine and interleukins

## Discussion

Vaccine development efforts for EV71 have mostly emphasized on inactivated whole-virus vaccines [Foo D et al., 2007], VLP vaccines (Podin Y et al., 2006), live attenuated vaccines [Arita M et al., 2007], polypeptide vaccines (Ooi EE et al., 2002), and DNA vaccines (Tu PV et al., 2007). Research into developing a vaccine against EV71 has gained a lot of interest and made remarkable advances over the past decade. Inactivated whole virus vaccines have been a main goal owing to their mature production technique and high levels of immunogenicity, safety, and stability, as indicated by a survey of the literatures published since 2007 on the development of EV71 vaccines. While this is the first global effort to systematically create EV71 vaccines, fast progress has been made with inactivated vaccines. Nevertheless, there are still a number of obstacles that must be overcome. During the last decade, Enterovirus 71 has shown a stronger tendency for mutation, and many new genotypes have since emerged and spread over the world (Bible JM et al., 2007; Chan YF et al., 2009; van der Sanden S et al., 2009).

For this purpose, Wu et al. tested the immunogenicity of inactivated whole-virus candidates grown in serum-containing and serum-free conditions in microcarriers and adjuvanted with alum (Wu SC et al., 2004). The immunogenicity of both candidates was high. Low levels of serum neutralizing antibodies were generated by oral vaccination with VP1 antigens, as reported by Chen et al. (Table 5), suggesting that oral immunization with VP1 antigens may need unique adjuvants to increase immunogenicity (Chen HF et al., 2006). Heat-inactivated whole-virus antigen and VP1 synthetic peptides (15 amino acid residues) were tested for their immunogenicity, and the results revealed that the VP1 synthetic peptides were less effective at eliciting an immune response (Foo DG et al., 2007). According to the results of Tung et al., analysis of VP1 DNA immunogenicity, the VP1 DNA vaccine candidate might generate serum neutralizing antibodies in a high dose level (100 g) in mice, which may not be practical in humans (Tung WS et al., 2007). While recombinant VP1 and DNA vaccines may be less expensive to produce, inactivated whole-virus vaccines seem to be more immunogenic overall. Vaccination is an effective, low-cost method of preventing disease spread across the worldwide (Murray, K.A et al., 2015). The immunoinformatics technique has been applied on a worldwide scale to lessen the financial and time requirements typically associated with the development process.

**Table 5.**
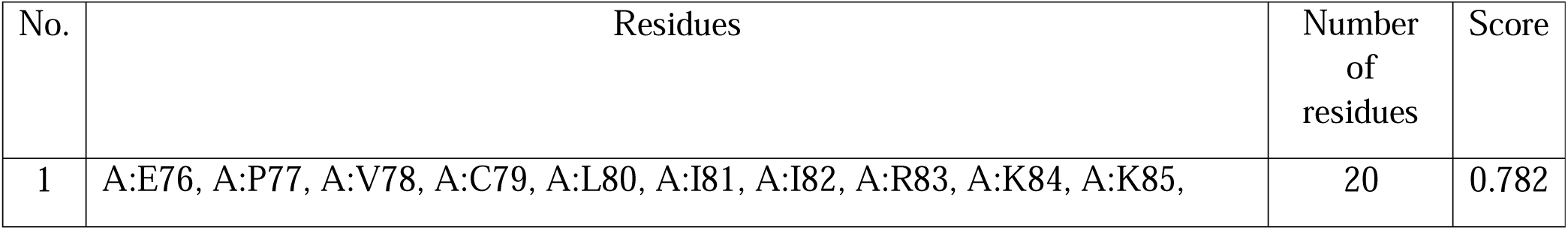
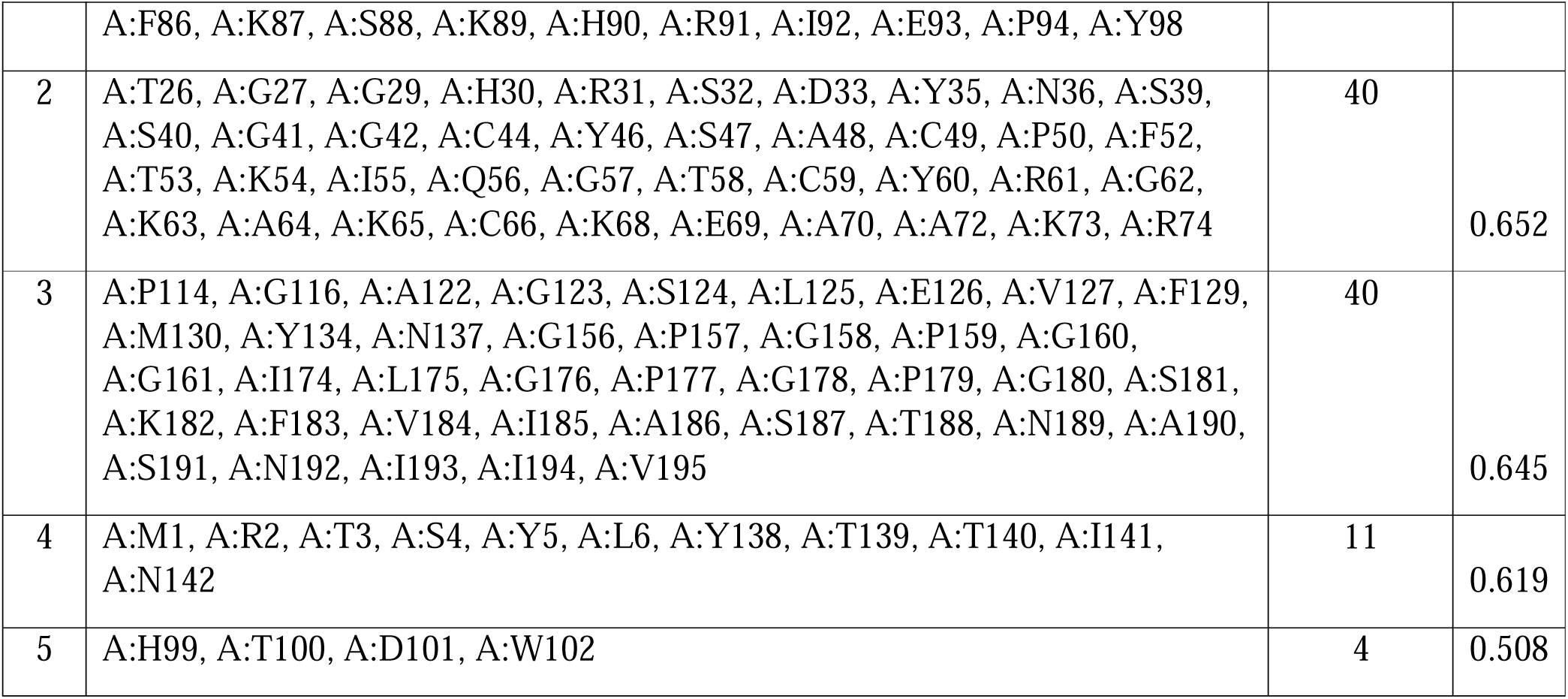
Confirmational discontinuous B-cell epitope

**Table 6.** Hydrogen bond (distance Å) and interaction of vaccine candidate with TLR3

| **Toll Like receptor-3** | **Vaccine** | **Distance (Å)** |
| --- | --- | --- |
| SER38 | THR3 | 1.8 |
| GLU33 | LYS85 | 1.7 |
| GLU33 | TRP102 | 1.9 |
| THR54 | ASP101 | 2.0 |
| SER132 | ILE194 | 1.9 |
| GLN107 | ASN137 | 2.0 |
| GLN107 | THR139 | 1.9 |
| ASN105 | THR139 | 2.0 |
| LYS102 | TYR146 | 1.6 |
| GLU127 | ARG169 | 1.8 |
| ASP153 | ARG169 | 1.9 |
| LYS201 | ASP170 | 1.7 |
| GLU175 | ARG171 | 1.9 |
| LYS200 | SER47 | 1.7 |

New approaches to vaccine formulation and development are being discovered at the genomics and proteomics level of pathogens (Khatoon, N et al., 2018; Sharma, A et al., 2021). Using immunoinformatics, multi-epitope-based vaccinations are a cutting-edge method for eliciting a targeted immune response while preventing reactions to additional, potentially harmful epitope antigens (Moise L et al., 2015). Better safety, the possibility of rationally engineering the epitopes for improved potency, and the capacity to concentrate immune responses on preserved epitopes are further possible benefits of multi-epitope-based vaccinations (Zhou W et al., 2015). In this investigation, we used a variety of bioinformatics instruments to design and assess a multi-epitope-based vaccination candidate against EV71. Moreover, immunological simulation was used to study the body’s reaction to the vaccine’s construction. In order to be used in a vaccine, a protein must be able to be presented on the surface and recognized by the immune system (Yasmin, T et al., 2016). These criteria, together with antigenicity and allergenicity analyses using VaxiJen and Aller-TOP v2, were used to determine the nucleocapsid protein that plays a major role in pathogenicity, as has been shown in earlier research (Parvizpour, S et al., 2009). In addition, the present research demonstrates that combining B-cell and T-cell epitopes to produce a robust and long-lasting immune response is an effective strategy for establishing cellular and humoral immunity.

## Declaration

### Conflict of Interest

The authors declare no known conflict of interest.

### Author contribution

## Acknowledgement

Research containing human participants or animals

## Table Legends

**Table-1** List of CTL epitopes predicted from EV-71 polyprotein to design multi epitope vaccine with immunogenic properties.

**Table-2** Prediction of HTL epitopes for multi epitope vaccine construct of EV-71 with immunogenic properties.

**Table-3** B-cell epitopes for multi epitope-based vaccine development.

**Table-4** Physicochemical properties of multi epitope vaccine construct.

**Table-5** Prediction of linear B-cell epitopes

**Table-6** List of hydrogen bond interaction between vaccine construct and TLR-3

## Notes

### Competing Interest Statement

The authors have declared no competing interest.

